# Efficient and precise discovery of LTR-retrotransposon families using HiRepEat

**DOI:** 10.64898/2026.10.14.759201

**Authors:** Anant Maheshwari, Eugene W. Myers, Pio Sierra, Vertebrate Genomes Project Consortium Phase I, Mara K. N. Lawniczak, Richard Durbin

## Abstract

Long Terminal Repeat Retrotransposons (LTR-RTs) are mobile genetic elements that replicate within host genomes using an RNA intermediate. Recent high quality genome assemblies provide new opportunities to study these repetitive elements across evolutionary timescales. We adapt the Myers *O*(*Nd*) *wave-front* alignment algorithm to produce a new fast LTR detection, clustering and phylogenetically-informed consensus approach for identifying LTR-RT families in new genome sequences. We present an iterative approach to identifying and removing repeats, initially tandem repeats, that we call TACO (TAndem COmpressor), and a corresponding coordinate mapping system we call SALSA (Sorted Array for Liftover of Sequence Addresses). Together these allow us to read and write in reduced genome space and search for LTR-RTs and tandem repeats in a series of operations. This greatly reduces confounding between different repeat types and allows us to uncover nested insertions. We show results from our clustering and consensus approach across the ***>***500 genomes in the Vertebrate Genomes Project, with analysis completing in most cases in under 60 minutes with 15Gb memory and 16 cores. We also present cross-genome LTR-RT clustering across 30 *Drosophila* species, as well as across the VGP, providing fine-grained resolution on species-sharing patterns of LTR-RTs and revealing candidates for horizontal transfer.

## 1 Background

Long Terminal Repeat Retrotransposons (LTR-RTs) are Class I transposable elements (TEs) that mobilize using an RNA intermediate. They typically encompass anywhere from 10-75% of eukaryotic genomes [1]. LTR-RTs are characterized by flanking same-orientation long terminal repeats (LTRs) that range from *∼*100bp to over *∼*5kb in size, with an internal sequence ranging from *∼*1kb to *∼*15kb. Functional autonomous LTR-RT elements contain open reading frames (ORFs) for protein-coding genes including gag (viral-like capsid) and pol (Protease, Reverse Transcriptase, RNaseH and Integrase). Closely-related Endogenous Retroviruses have an additional ENV domain, which encodes for envelope proteins enabling their extracellular movement via viral particles [2].

Existing widely used repeat detection frameworks such as RepeatModeler2 [3] and EDTA [4] internally leverage tools such as LTR Finder [5] and LTRharvest [6] for structural detection, followed by LTR retriever [7] for post-processing and false positive removal. Polymorphism-based tools for TE detection, such as pantera [8] and Pannagram [9] provide an orthogonal approach based on comparing genomes and clustering structural variants into putative families, followed by classification. A variety of LTR specialist tools have been proposed, for example Look4LTRs [10].

After extracting putative LTR-RTs from an input genome, these tools generally rely on CD-HIT [11] to cluster instances together, followed by MAFFT [12] for deriving a consensus sequence from each cluster, to define putative LTR-RT families. Families are typically defined following Wicker et al. [1] using the 80/80/80 rule, that is 80% of the sequence must align to 80% identity over at least 80bp, and CD-HIT clustering thresholds are set accordingly. However, the CD-HIT clustering approach relies on greedy heuristics based on matching members to a representative sequence, which does not guarantee an optimal solution, and furthermore the clusters formed can be loosely connected due to transitive alignment. This can lead to lossy and unrepresentative consensus sequences over loosely aligned structures, where cluster members can diverge from each other much further than the 80/80/80 rule definition.

Furthermore, LTR-RTs frequently occur in nested structures involving both tandem repeats and other LTR-RTs, inserting into each other, which confounds these widely used detection methods based on naive clustering and limits the detection of older TE families. For example, if an LTR-RT instance is nested inside another LTR-RT instance e.g. a younger insertion inside an older insertion, using a naive CD-HIT clustering approach the outer LTR-RT instance would never cluster with family members due to the internal insertion confounding the alignments. Towards handling this nested and confounding repetitive content, various advances have been made in the TE annotation field. For example, TE-greedy-nester [13] uses a greedy recursive algorithm for nested TE recovery; however it also runs LTR Finder [5] (as used by EDTA, RepeatModeler2, etc.) as a backend, and does not report libraries but rather nesting order and insertion ages.

The *O*(*Nd*) *wavefront* alignment algorithm [14] [15] [16] provides a fast guaranteed optimal alignment given maximal divergence rate *d*, by leveraging greedy extension operations in order to avoid exploring the entire dynamic programming matrix. Here we leverage this and other algorithms to achieve an efficient and precise approach to LTR detection, subsequent clustering, and phylogenetically-informed consensus building to build candidate families. We also describe TACO (**TA**ndem **CO**mpressor) and SALSA (SALSA: **S**orted **A**rray for **L**iftover of **S**equence **A**ddresses), which we use to delete high-confidence detected tandem repeat and LTR-RT instances from the genome, writing out a compressed genome sequence together with the coordinate-mapping information needed to reverse the process after analysis of the reduced genome. Overall, the described approach below forms an end-to-end system that takes as input a genome and outputs consensus sequences for the LTR-RT families discovered, together with, optionally, a multiple alignment for each family of full length representatives.

We apply this pipeline to 577 genome sequences from the Vertebrate Genomes Project (VGP) phase I dataset [17] and 30 *Drosophila* genomes [18], and show bench-marking results demonstrating high precision and computational efficiency, together with an overview of LTR-RT distribution and properties across vertebrates.

## 2 Methods

### 2.1 Iterative tandem repeat collapsing with TACO/SALSA

Tandem repeats are a major contributor to genomic repetitive content that can confound the identification of LTR direct repeats LTR-RTs, and thus become sources of false positives for LTR-RT detection. Furthermore, these sequences are particularly difficult to align through as there are many possible alignments to explore over dense tandem-repeated regions, for instance see the dense tandem array in Supplementary Information Figure 1. Based on this intuition, we take a preprocessing approach that significantly accelerates the downstream TE detection tasks. FasTAN (Myers, https://github.com/thegenemyers/FASTAN) is a tandem repeat detector that detects tandem arrays with repeat lengths between 2bp to 30kbp, allowing for up to 30% divergence. We iteratively use FasTAN to detect tandem repeats, and then compress each tandem repeat down to just the first copy using TACO (**TA**ndem **CO**mpressor) as shown in Figure 1a(i). Multiple rounds of FasTAN + TACO are iterated until no more tandem repeats are found. Each round uncovers higher-order tandem repeats, for example leading to the compression of a 35kb tandem repeat region in *Anopheles funestus* to be compressed down to just 286bp as shown in Supplementary Information Figure 1, and Supplementary Information Algorithm 1.

**Fig. 1:**
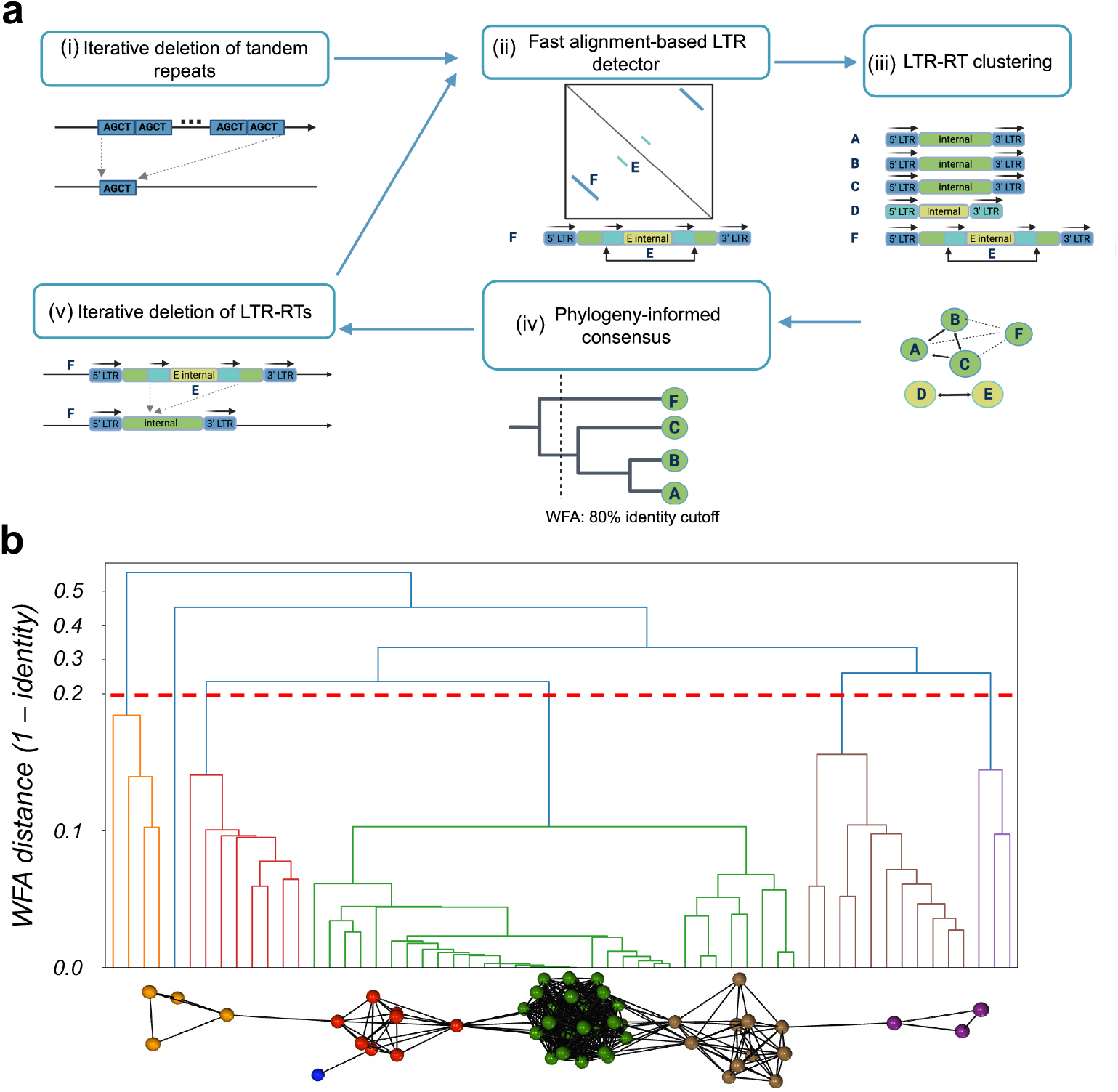
End-to-end LTR-RT detection and analysis approach. **a** i) iterative detection of tandem repeats with FASTAN + TACO/SALSA collapse, preserving the first copy of each tandem repeat to collapse hierarchical tandem structures. ii) detection of a set of candidate LTR-RTs flanked by direct terminal repeats using FastLTR. iii) candidate clustering: nodes are candidates, with solid edges joining them if both the terminal repeats and internal sequences satisfy the 80/80/80 rule. iv) phylogenetic consensus generation for each cluster (see **b**). v) collapse sequences A, B, C from the genome, and also collapse sequences D and E. On the next iteration the reduced F will correctly cluster with A, B, C. **b** A real cluster (from *Astatotilapia calliptera*) with edges between nodes if both their terminal repeats and their internal sequences align to 80% identity, showing transitive connectivity. The reconstructed phylogeny shows sequence divergence on the y-axis and our threshold-based split into five families and a singleton.

This preserves the unique sequence content of the tandem repeat since we still maintain one copy. We manage the associated coordinate transformations in SALSA (**S**orted **A**rray for **L**iftover of **S**equence **A**ddresses), which uses binary search for fast coordinate transformations, allowing us to seamlessly compress and decompress. The implementation writes two ONEcode files [19] with suffixes .1seq and .1taco at each TACO iteration, where the .1seq file stores the sequence with removed tandem repeat copies, and the .1taco file stores each liftover event i.e. coordinate mappings per compression operation, alongside the removed DNA sequence such that this compression is lossless. The resulting TACO-compressed genomes (.1seq) are used for downstream LTR-RT detection, preserving the ability to map sequence coordinates back via SALSA.

### 2.2 LTR-RT candidate detection

Identification of LTR-RT candidates works by searching first for clusters of 8-mers recurring at a consistent spacing in sliding 32kbp windows with 12kbp overlap, and then performing fast *O*(*Nd*) alignments to verify matches, using program FastLTR. This process will report candidate segments with direct (non-inverted) terminal repeats that are up to 30% divergent, in spacing ranges between 2kbp and 20kbp, which is the expected range for full-length LTR-RTs. Thus, by running this on an input genome, we have a set of structural candidate LTR-RTs (example shown in Figure 1a(ii)). However many of these will still not be LTR-RTs but rather various other forms of confounding repetitive content, despite pre-removal of much of this confounding content by our FasTAN/TACO process.

### 2.3 LTR-RT clustering

To look for dispersed families of LTR-structured sequences that are more likely to be LTR-RT TEs, we next take all of our putative LTR-RT sequences and construct clusters using the following approach as shown in Figure 1a(iii), and Supplementary Information Algorithm 2, 3, 4. We first compare just the terminal repeats, considering both forward and reverse complement orientations. We compute canonical 12-mers over input candidate instances, and restrict to pairs sharing at least 3 of these canonical 12-mers. We then choose an alignment anchor centered around the most dense region of 12-mer matches, and run *O*(*Nd*) *wavefront* alignment extending from both directions of the anchor, and requiring less than 20% divergence in both directions. Given that the terminal repeats match to at least 80% identity, we confirm alignment of the internal sequence between the repeats by following the same seed, anchor, and WFA alignment process. We take matches between candidates that pass both the terminal repeat and internal alignment checks as passing the 80/80/80 family definition. We require a minimum of three LTR-RT instances to form a cluster, at a minimum of 2kbp length, so HiRepEat is not suited to detecting low-copy, fragmented LTR-RTs and is aimed towards high-confidence full-length detection.

### 2.4 Phylogeny-informed consensus approach

After computing connected components, we have clusters (components) from which we wish to derive consensus sequences. Consensus sequences can be thought of as a representative/ancestral sequence of the putative TE family, which can be used for repeat masking or comparative studies of TE families. However, there is a transitive issue as alluded to earlier, where within a connected component under the *80/80/80* rule we may have A connected to B and B connected to C, but A not connected to C, as illustrated for example in Figure 1b, where multiple families are transitively connected. To resolve this issue, we build a tree of the instances in a cluster using Supplentary Information Algorithm 5. We leverage our pre-computed WFA similarity measures from our clustering algorithm to build a guide tree via UPGMA, filling in missing similarities based on shortest paths within the connected component. We then cut the UPGMA tree at 0.20 identity threshold such that we ensure our clusters are internally following the 80/80/80 rule, and proceed with progressive alignment within each subtree based on *O*(*Nd*) *wavefront* alignment of consensus sequences. At each step, if the two ingroup bases at some consensus position conflict then we tie-break using a closest outgroup sequence. This allows us to derive meaningful consensi based on the underlying LTR-RT family phylogeny (Figure 1a(iv)).

#### 2.4.1 Classifying LTR-RTs and iterative detection using TACO/SALSA

After we detect, cluster, and derive consensus sequences, we next classify LTR-RT families using TEsorter [20], which leverages the REXdb database [21] of both plant and metazoan LTR-RT protein domains, alongside the GyDB database [22]. We remark that our framework is extensible and other classifiers/databases of choice can be adapted for this step.

We then proceed to use our TACO/SALSA infrastructure to delete all instances of our high-confidence LTR-RT detected families from the genome, as shown in Supplementary Information Algorithm 6 and Figure 1a(v). This emits new .1seq and .1taco files, similar to the tandem repeat case, but deleting entire repeats as opposed to storing the first copy. We next apply the FastLTR detection again to the reduced genome, followed by consensus generation combining with the previously detected candidates, and then classification and another round of TACO/SALSA deletion of LTR-RT instances. We repeat this whole process until convergence (Figure 1). This progressive deletion approach, eating up copies of validated repeats, can be thought of as rewinding TE insertions by deleting them in rounds, and finds nested insertions that are otherwise invisible to standard clustering methods.

## 3 Results

We implemented the LTR-RT family discovery and classification process as software package **Hi**erarchical **Rep**eat **Eat**er or HiRepEat (named to describe the principle of hierarchically “eating” up repeats), as described in the Methods section.

### 3.1 Benchmarking HiRepEat LTR-RT detection

We first compared HiRepEat to curated annotations and previous LTR-RT discovery tools, for three test species *O. sativa* (rice), *T. guttata* (zebrafinch), *G. gorilla* (gorilla). To do this we ran HiRepEat, LTR retriever [7], Look4LTRs [10] and pantera [8] on each species genome, and for HiRepEat, pantera and LTR retriever we then used RepeatMasker [23] to mask the respective genomes (Look4LTRs annotates the locations of found LTR families automatically as part of its standard output). We chose LTR retriever and Look4LTRs for comparison because LTR retriever, which applies both LTRharvest [6] and LTR FINDER [5], is a standard LTR-RT discovery tool used for example in both RepeatModeler2 [3] and EDTA [4], whereas Look4LTRs is a recent efficient tool that searches explicitly for nested insertions. We added pantera as an alternative, polymorphism-based approach that gives high quality full length consensus sequences for recently mobile families, so has high precision/specificity, though with lower recall/sensitivity.

We describe the information flow for HiRepEat in Supplementary Table 2 on rice, zebrafinch, gorilla. For example, on zebrafinch (1,141 Mb genome) we see that HiRe-pEat goes through 8 FasTAN TACO/SALSA iterations, removing a total of 81 Mb of tandem repeats prior to LTR-RT steps. Over the course of 4 LTR-RT rounds, FastLTR detected 15,320 candidate LTR instances, of which 131,573 pairs pass the terminal repeat matching, followed by 28,422 pairs passing the internal sequence matching. After filtering out 480 edges due to them sharing flanking sequences, this results in a graph with 5,861 nodes and 27,942 edges. This process results in 75 connected components with *>* 3 nodes, upon which we apply our phylogeny-based consensus splitting approach resulting in the construction of 125 consensus sequences, out of which 99 are classified as LTR-RT families based on protein domain evidence.

Precision, recall, F1 along with runtime and memory information are provided in Figure 2a. Comparing the LTR-annotated bases against the curated set for zebrafinch, we see that HiRepEat achieves a precision of 0.91 and recall of 0.71, achieving the highest F1 score of 0.79. LTR retriever achieves the highest recall (0.89) but at very low precision (0.44), interestingly calling 115Mb of the genome to be LTR-RT despite the curated set only indicating 57Mb. Pantera achieves the highest precision (0.93) but much lower recall (0.39) which can be attributed to it only being able to detect polymorphic insertions. In rice, we see a similar pattern where HiRepEat achieves a high precision/F1 relative to other LTR-specialist callers while trading off some recall, with a significant runtime acceleration and requiring less memory. Figure 2b shows the intersections of overlapping LTR-RT Mb detected by the 4 tools, for each of the 3 genomes.

**Fig. 2:**
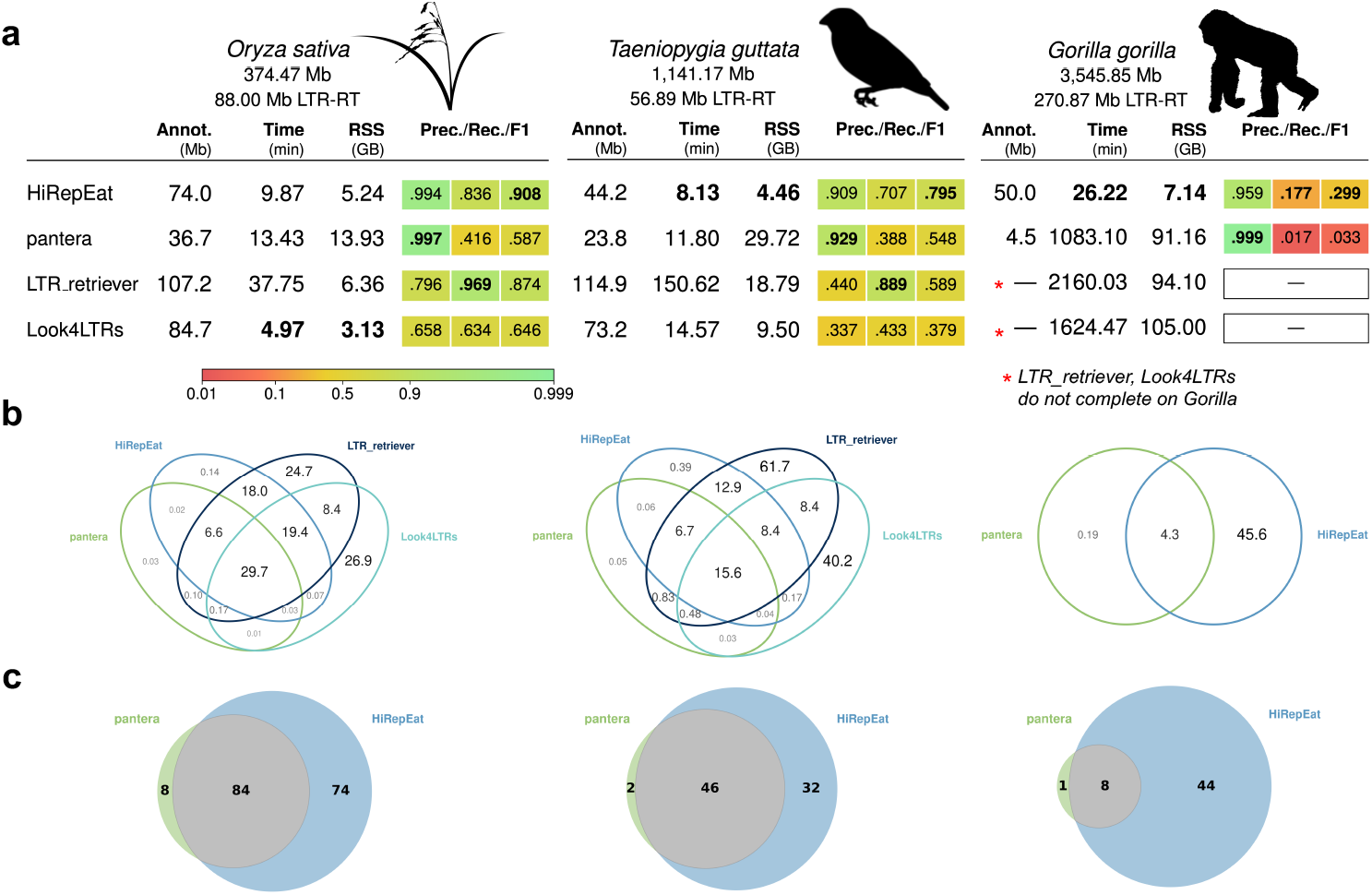
Benchmarking on *Oryza sativa* (rice), *Taeniopygia guttata* (zebrafinch), *Gorilla gorilla* (gorilla). All experiments were carried out with 16 cores, 105Gb max memory, with a time limit of 36 hours. On gorilla, LTR retriever timed out after 36hr, and Look4LTRs ran out of memory at 105Gb after 27:04hr spent processing chromosome 1, so only pantera and HiRepEat results are shown. **a**) Comparing LTR-RT genome annotations of each tool against the expert-curated set, to measure annotated Mb, precision, recall, F1 as well as runtime, and max memory usage. **b**) Showing overlap of masked genome (in Mb) by pantera, HiRepEat, LTR retriever, Look4LTRs. Note, size of venn overlaps do not reflect intersection sizes. **c**) Benchmarking at family-level with pantera using blastn (80% pident and 80% qcovs) to measure concordance of families found; grey indicates families that match.

We remark that on gorilla, HiRepEat runs in *∼*26 minutes, whereas pantera runs in 18 hours, and LTR retriever times out after 36 hours. Although Look4LTRs is the fastest method on the smallest genome (*Oryza sativa*), on *Gorilla gorilla* it goes above max memory (105Gb) after processing chromosome 1 for 27 hours. We remark that even though HiRepEat displays a high precision on gorilla (0.96), it has a significantly lower recall (0.18) than for zebrafinch and rice, despite being higher than pantera’s recall (0.017). This gap can be explained by the presence of ancient fragmented non-autonomous endogenous retroviruses (ERVs) with no intact proteins present (such as families of ERVL-MaLR), which do not have long terminal repeats, and are thus by construction inaccessible to HiRepEat as well as to other structural LTR-RT callers. We report this as a limitation of our method, and recommend using HiRepEat for biological questions requiring precise detection of full-length LTR-RTs.

We next compared the families generated by HiRepEat against LTR-RT families found by pantera [8], by comparing the consensus sequences with blastn with pident *≥* 80% and qcovs *≥* 80% in both directions. Pantera uses an orthogonal detection strategy based on polymorphism between haplotypes to find recent insertions, which tends to accurately identify full length sequences for recently active TEs. We would hope that HiRepEat will find all the LTR-RTs that pantera finds, in addition to recovering families represented only by older non-polymorphic insertions that pantera does not see. The results in Figure 2c show this to be the case, with HiRepEat recovering almost all of the LTR-RT families detected by pantera. Of the missed cases, the majority have consensus sequences that are 69-80% identical to some HiRepEat consensus sequence thus fail the 80% identity threshold. Additionally, we see three missed cases of ERV families with no intact terminal repeat or no protein matches (non-autonomous).

Overall, we see that HiRepEat achieves a high precision and good recall comparable to the state of the art tools, while running an order of magnitude faster than those tools. This enables scaling to large numbers of genomes and further to particularly large, unexplored genomes. We explored this scaling next.

### 3.2 Studying the evolution of LTR-RTs across vertebrates

We ran HiRepEat on the 577 chromosomal genome assemblies from the Vertebrate Genomes Project Phase I release [17], finding 130,110 LTR-RT families. Figure 3a) shows how runtime, memory and number of families discovered scale according to genome size, with the majority running in under 60 minutes within 15GB memory, using 16 cores. We note that HiRepEat runs successfully on *>*25Gb amphibian genomes, for example finding 8,119 families in 23.6 hours and 50.9GB memory for *A. mexicanum* (axolotl) and 7,101 families in 17.4 hours and 96.7GB memory for *T. cristatus* (Northern crested newt).

**Fig. 3:**
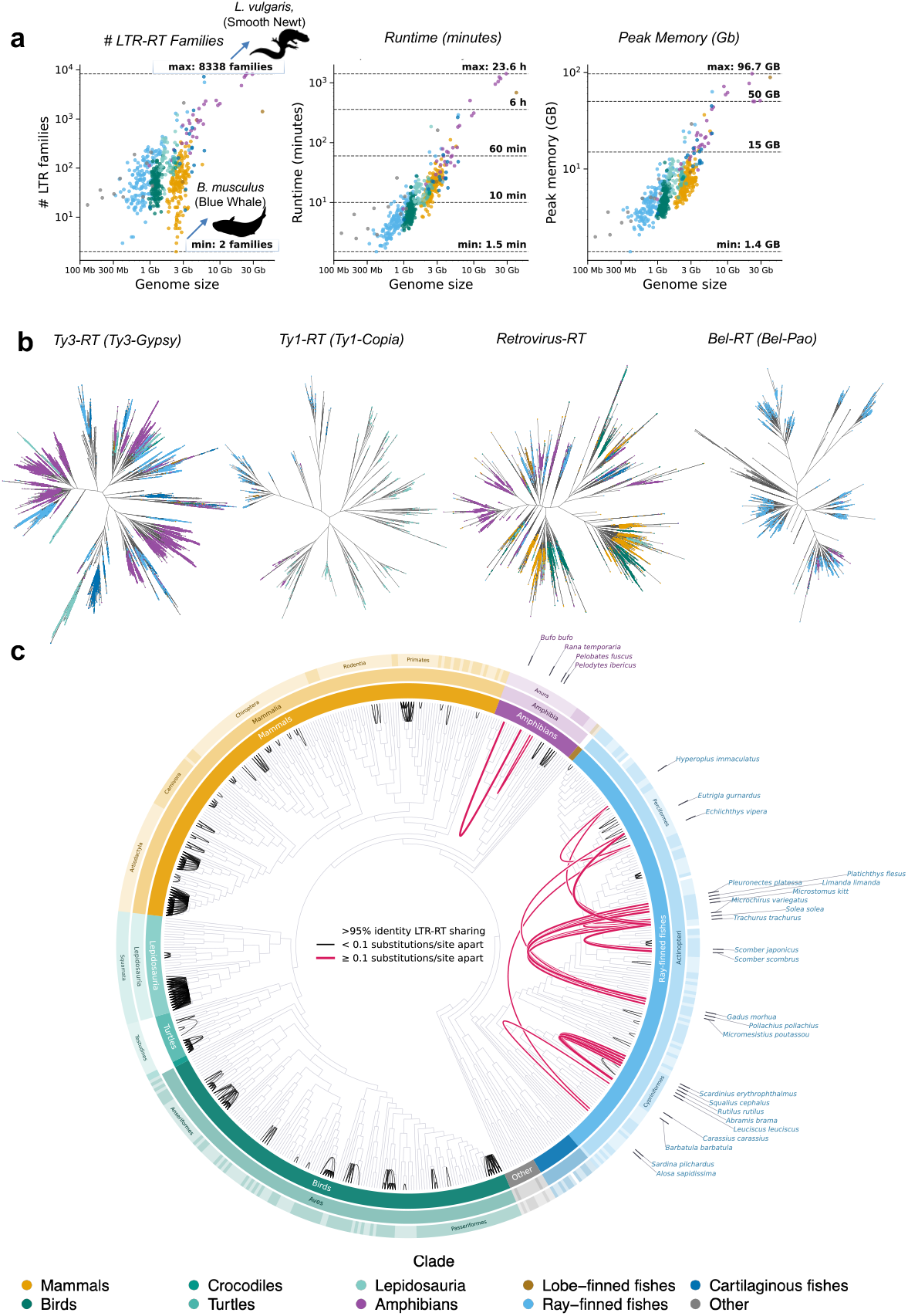
Running HiRepEat on 577 genomes from the Vertebrate Genomes Project. **a)** Scaling properties of runtime, memory usage, and # LTR-RT families detected across the vertebrates, showing genome size dependencies. **b)** Reverse-transcriptase protein-based phylogeny, using 105,437 reverse transcriptase domains from HiRepEat LTR-RT consensus sequences across the 577 vertebrate genomes. **c)** Clustering LTR-RTs discovered by HiRepEat across the VGP, at 95% identity, imposed on top of the VGP phase I ROADIES species tree. Highlighted *>* 95% identity LTR-RT cross-species edges that connect species differing by more than 0.1 substitutions per site (pink). Distant-range sharing patterns suggest potential horizontal transfer candidates, particularly in fish.

Beyond their DNA consensus sequences, the families identified generate a dataset of protein domains, including 105,437 reverse transcriptase (RT) domains. These can be used for subclassification of LTR-RT superfamilies, as in the REXdb database of plant transposable element protein domains [21]. We used FastTree [24] to infer approximately-maximum likelihood trees for superfamilies Ty3/Gypsy, Retrovirus, Bel-Pao, Ty1/Copia RT domains (median amino acid lengths 173, 203, 222, 255 respectively; Figure 3b). Ty3/Gypsy are primarily represented by amphibians and fishes, with minimal bird and mammal sequences. Retroviruses, in contrast, are dominated by birds and mammals. In vertebrates Ty1/Copia has previously been reported to be largely restricted to non-amniotes, retained in fishes, amphibians, reptiles but lost in birds and mammals [25], and we indeed recover Copia RT domains predominantly from fishes, amphibians, and reptiles. We additionally see Bel/Pao elements abundant in fishes and the VGP outgroup invertebrates but essentially lost in other clades. We interestingly see some cross-clade sharing patterns, for example birds and mammal RT domains appear to mix together and not display a clear clade-like split, perhaps hinting at ancient LTR-RT families that were shared across the clades or horizontal transfer events.

### 3.3 Cross-genome clustering mode reveals horizontal transfer candidates

Motivated by the suggestion of clade switching in the RT trees, we next asked questions around broader LTR-RT sharing patterns across the vertebrates. For each of the 130,110 HiRepEat consensus sequence across all the vertebrate species, we re-aligned cluster members back to the consensus sequences, to choose a representative sequence closest to the consensus. We then performed our LTR-RT clustering algorithm over these representative sequences using HiRepEat, requiring 95% identity (instead of 80%) for both the LTRs and the internal sequence in order to form edges in a graph, where nodes are representative sequences. This process resulted in 3,446 cross-species edges, which we display in Figure 3c), imposed on top of the ROADIES [26] v0.0.0 The Vertebrate Genomes Project Phase I species tree [17].

Notably, we see several patterns of long-range sequence sharing that cross large phylogenetic distances, particularly in fish. For example we have a *>* 95% identity Bel/Pao element found in blue whiting (*Micromesistius poutassou*), Atlantic horse mackerel (*Trachurus trachurus*), thickback sole (*Microchirus variegatus*), European flounder (*Platichthys flesus*) and European plaice (*Pleuronectes platessa*). These species are from three different orders, with most recent common ancestor well over 100 million years ago, but overlap geographically in the Northeast Atlantic, which could facilitate horizontal transfer.

To further explore such cases, we use the ROADIES tree pre-computed branch lengths (substitutions/site), allowing us to highlight horizontal transfer (HT) candidates that connect particularly divergent species (more than 0.1 substitutions/site, yet sharing *>* 95% identity LTR-RT insertions). All but two of these candidates are found in teleost fish, consistent with previous observations [27] [28]. For example, in the case described above we have blue whiting has *∼*0.5 subs/site based on the ROADIES tree with respect to the others. The other two examples are between pairs of amphibians, the common toad *Bufo bufo* and common frog *Rana temporaria*, and the common spadefoot toad *Pelobates fuscus* and Iberian parsley frog *Pelodytes ibericus*, all of which lay external eggs in water, consistent with proposals that LTR-RTs may transfer between species during external aquatic fertilisation [29]. The automated generation of these candidates show the benefit of our scalable methods for studying LTR-RT biology.

A powerful extension of HiRepEat is the cross-genome mode, due to our fast allagainst-all clustering approach. This provides a means to directly cluster LTR-RTs across multiple genomes at once, such that we can see the sharing patterns at the resolution of individual TE instances, as we use individual LTR-RT instances rather than representative consensus sequences. We demonstrate this by directly clustering all the LTR-RT instances across 30 *Drosophila* species chromosomal assemblies from Gebert et al. [18], as shown in Figure 4. This process took 2 hours using 16 cores, with a max memory of 16.5Gb. Visualizing the clusters formed (coloured by species) we see a large diversity of densities and topologies of TE family clusters, with sharing patterns primarily driven by within or nearby species. However, there are also some notable off-diagonal cases, such as *D. sechellia* sharing an *LTR/Gypsy* family to *∼*99% sequence identity with *D. willistoni* even though these species are thought to have diverged *∼*50 million years ago. These off-diagonal cases suggest candidate horizontal transfer cases. We note that related LTR-RT horizontal transfer cases involving *D. willistoni* have been reported in the literature and are speculated to be due to recent human activity influencing geographical patterns [30] [31].

**Fig. 4:**
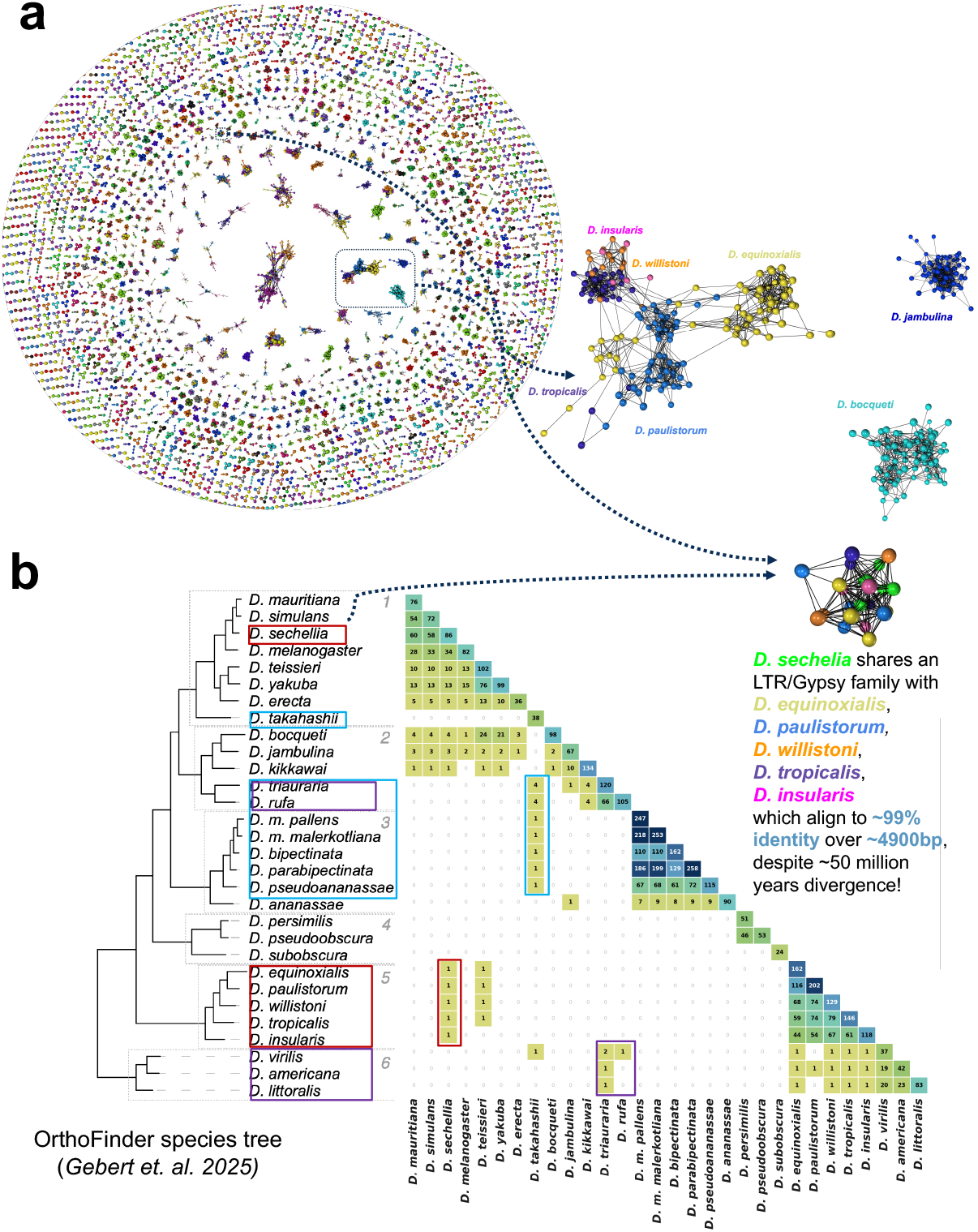
Cross-genome mode clustering 30 *Drosophila* Species from Gebert et al. (2025). **a**) Real clusters formed where every node is an LTR-RT instance (in any of the 30 *Drosophila* genomes), edges connect if they align 80/80/80, and colours indicate species. **b**) Bottom heatmap shows the number of shared LTR-RT families i.e. clusters with members from each species, with the Gebert et al. Orthofinder-based species tree shown on the left. Sharing patterns between distant species indicate potential horizontal transfer cases.

## 4 Discussion

We have introduced a new computational method HiRepEat for efficient, precise discovery of full-length LTR-RT families from genome sequences, within and between species. A key feature of our approach is to iteratively “eat” up previously identified repeat instances, starting with tandem repeats found with FasTAN. This avoids consideration of many of the false positive candidates generated by other LTR-RT detection systems, which can report tandemly copied sequences as candidate LTRs. This reduction in candidate generation both leads to higher precision and substantially greater speed. Subsequent rounds of “eating” already found LTR-RT family instances support the correct discovery of nested LTR insertions.

We remark that due to our clustering thresholds requiring a minimum of three copies, alongside our tandem repeat removal process, HiRepEat will not find all forms of fragmented and degraded LTR-RTs described in the literature, such as ancient endogenous retroviruses [32] and tandem LTR-RTs [33]. We also note that some Class I superfamilies are not detectable within our current framework, namely LTR/DIRS and LTR/Ngaro, since these have split/inverted repeats and are by construction not visible to HiRepEat. Nevertheless, our accelerated runtime and higher precision enables capturing the major classes of LTR-RTs efficiently and sensitively, which is a major hurdle for existing TE detection frameworks. For example, we are able to analyse gorilla in *∼* 26 minutes, contrasting to pantera running in 18 hours and other methods not completing, and large genome amphibans such as axolotl, which have provided major challenges for previous TE detection approaches, within a day.

We also remark that our HiRepEat framework and iterative detection approach is extensible to other classes of repeats beyond tandem repeats and LTR-RTs. We anticipate releasing future versions that include DNA transposons for which candidates can be found as containing terminal inverted repeats, and other classes using appropriate specialised detection systems. Further, we suggest that the compressed genomes that we end up with may be preferable for some downstream analyses to repeat-masked genomes, that are naturally fragmented by the inserted repetitive sequences. Our SALSA framework will support the necessary coordinate transformations to convert back from locations in the compressed sequence space to the original coordinates.

## 5 Conclusions

HiRepEat provides a new efficient package for iterative detection of LTR-RT candidates using FastLTR, clustering and phylogenetically-informed consensus building, and collapse of discovered repeats using TACO/SALSA, together forming an end-to-end system for LTR-RT analysis. We apply this to all the genomes in the Vertebrate Genomes Project phase I release, demonstrating to ability to scale automated detection and analysis across the ever-increasing number of high-quality genome assemblies becoming available. We further demonstrate cross-species clustering of LTR-RTs, both across 577 vertebrate species and across 30 *Drosophila* assemblies, revealing horizontal transfer candidates. This work enables efficient LTR-RT library construction, which is a significant bottleneck in existing transposable element detection and annotation methods [3] [4].

## Availability of data and materials

The code for HiRepEat is available at https://github.com/AnantMaheshwari/HiRepEat, which also includes a singularity image indicating all software library versions used. The curated set for *Oryza sativa* was downloaded from https://github.com/oushujun/EDTA/tree/master/database and the curated sets for *Taeniopygia guttata* and *Gorilla gorilla* were downloaded from https://tinyurl.com/HiRepEat-curated. The ROADIES v0.0.0 tree was downloaded from https://github.com/VGP/vgp-tree-release. All HiRepEat libraries generated and horizontal transfer candidates described are available on Zenodo (DOI 10.5281/zenodo.23372956).

## Acknowledgements

We would like to thank the members of the Vertebrate Genomes Project Repeats subgroup for stimulating engagement, and in particular Arian Smit for curating libraries for zebrafinch and gorilla. We also thank the Durbin, Karam-Teixeira and Lawniczak groups for useful discussions, and all those contributing samples and genome sequences towards the Earth Biogenome Project goals.

## Author contributions

AM developed the overall HiRepEat software framework, including the TACO/SALSA, clustering, phylogeny and consensus generation programs, carried out all the analyses and wrote the first draft of the manuscript. EWM developed the FasTAN and FastLTR programs. PS provided expertise in TE family detection and classification. The VGP provided genome sequences and related resources. MKNL and RD supervised the project. All authors revised and approved the manuscript.

