## Supplementary information for "Efficient and precise discovery of LTR-retrotransposon families using HiRepEat"

#### Figures

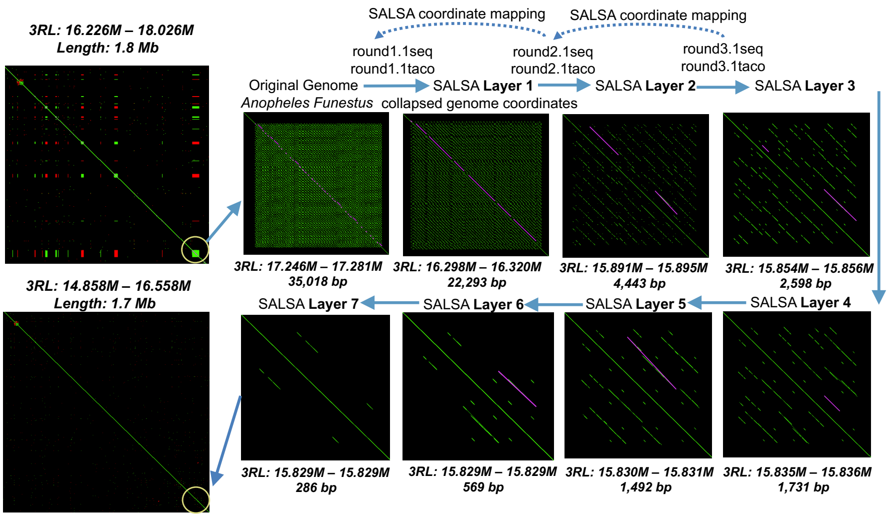


**Supplementary Figure S1**: TACO/SALSA tandem repeat collapsing of a 35Kb region in *Anopheles funestus*. This shows a self-alignment, pink lines are the tandem repeats detected by FasTAN at each round which we collapse to just the first copy, and green dots are based on a 19-mer dotplot (red dots are inverted). Through 7 compression operations (7 SALSA layers) the 35Kb region is compressed to 286bp.

### Algorithms

**Algorithm 1** Iterative Tandem Repeat Collapse (FasTAN + TACO/SALSA)

max_rounds = 10 (in principle, $\infty$)
**repeat**
  $R \leftarrow$ FasTAN(sequence)
  **for** each repeat $x \in R$ **do**
    Collapse $x$ to first copy via TACO
  **end for**
  New SALSA layer: `.1taco` + `.1seq`, decrement max_rounds
**until** $R = \emptyset$ or max_rounds reached

**Algorithm 2** K-mer Seeding and Anchor Selection

Compute canonical $k$-mers ($k = 12$) for all sequences
Group hits by pair $(i, j)$; keep pairs with $\geq$ min_shared = 3 matching canonical 12-mers
**for** each passing pair $(i, j)$ **do**
  Compute diagonals $d = \text{pos}_i - \text{pos}_j$ and anti-diagonals $ad = \text{pos}_i + \text{pos}_j$
  **if** $\text{var}(ad) < \text{var}(d)$ **then**
    is_rc $\leftarrow$ true
  **end if**
  Select anchor: hit closest to mean diagonal (or anti-diagonal if RC)
**end for**
**return** candidate pairs with anchors $[i, j, \text{anchor}_i, \text{anchor}_j, \text{is\_rc}]$

**Algorithm 3** Anchored $\mathcal{O}(Nd)$ *Wavefront* Alignment

**for** each candidate pair $(i, j, \text{anchor}_i, \text{anchor}_j)$ **do**
    max_edit $\leftarrow 0.2 \times \max(\text{len}_i, \text{len}_j)$
    Run WFA from anchor, extending left and right
    **if** required edits > max_edit **then**
        identity $\leftarrow -1$ (reject)
    **else**
        identity $\leftarrow 1 - \text{edit\_distance}/\max(\text{len}_i, \text{len}_j)$
    **end if**
**end for**
**return** $\{(i, j) : \text{identity}\}$ for passing pairs

**Algorithm 4** LTR-RT Clustering Graph Construction

Seed and anchor all LTR pairs (5', 3', RC) (**Algorithm 1**)
Run anchored WFA on LTR regions (**Algorithm 2**)
Flank filter: remove pairs $(i, j)$ where 100bp flanks also pass WFA, to remove tandem repeats
Seed and anchor internal regions (**Algorithm 1**)
Run anchored WFA on internal regions (**Algorithm 2**)
Retain edge $(i, j)$ only if it passes *both* LTR and internal alignment
Build graph $G$: nodes = element indices, edges $(i, j)$ defined by WFA identity
Find connected components of $G$; discard components with $<$ min_cluster_size $= 3$ members
**return** $G$ and retained clusters

**Algorithm 5** LTR-RT Progressive Alignment for Consensus Construction (UPGMA-style)

Build sparse distance matrix from pairwise identities, filling missing entries via shortest paths
Build UPGMA tree $T$ from $D$: $\text{dist}(AB, C) = \frac{|A| \cdot \text{dist}(A,C) + |B| \cdot \text{dist}(B,C)}{|A|+|B|}$
Cut $T$ at threshold 0.20 into subfamilies (split if merge distance $> 0.20$)
**for** each subfamily, process bottom-up over $T$ **do**
    **while** more than one profile remains **do**
        Take next sibling pair $(A, B)$ from UPGMA order
        Compute consensus strings $\text{cons}_A$, $\text{cons}_B$
        Anchor-seed and run WFA on $\text{cons}_A$ vs $\text{cons}_B$ (edit budget $= 0.20 \times \max(\text{len}_A, \text{len}_B)$)
        **if** outgroup $C$ available and ambiguous columns exist (top_count $< 1.5\times$ second_count) **then**
            Align $\text{cons}_{AB}$ vs $\text{cons}_C$; use $C$ to disambiguate tied columns
        **end if**
    **end while**
    **output** consensus sequence
**end for**

**Algorithm 6** Iterative LTR-RT Collapsing (TACO + SALSA)

max_rounds = 10 (in principle, $\infty$)
Initialize graph $G$
**repeat**
  $D \leftarrow$ FastLTR
  $d \leftarrow$ *TACO lift* $D$ to previous round coordinate space, find detections $d \in D$ that overlap last round's collapsed regions
  WFA new detections $d$ against all other nodes in $G$ (including previously collapsed LTR-RT instances); add corresponding new nodes $d$ and edges based on WFA 80% identity to $G$
  $C \leftarrow$ Connected Components($G$)
  Build consensus (**Algorithm 5**) and classify $C$
  **for** each new sequence belonging to a classified cluster **do**
    Collapse all LTR-RT instances corresponding to that cluster via TACO/SALSA
  **end for**
  New layer: `.ltaco` + `.lseq`, decrement max_rounds
**until** $D = \emptyset$ or max_rounds reached
